# Medial Orbitofrontal Cortex Regulates Instrumental Conditioned Punishment, but not Pavlovian Conditioned Fear

**DOI:** 10.1101/2020.05.12.092205

**Authors:** Cassandra Ma, Philip Jean-Richard-dit-Bressel, Stephanie Roughley, Bryce Vissel, Bernard W. Balleine, Simon Killcross, Laura A. Bradfield

## Abstract

Bidirectionally aberrant medial orbitofrontal cortical (mOFC) activity has been consistently linked with compulsion and compulsive disorders. Although rodent studies have established a causal link between mOFC excitation and compulsive-like actions, no such link has been made with mOFC inhibition. Here we use excitotoxic lesions of mOFC to investigate its role in sensitivity to punishment; a core characteristic of many compulsive disorders. In our first experiment, we demonstrated that mOFC lesions prevented instrumental conditioned punishment learning in a manner that could not be attributed to differences in Pavlovian conditioned fear. We then showed that increasing the frequency of punishing outcomes allowed mOFC-lesioned animals to overcome their initial deficit. Our second experiment demonstrated that the retrieval of instrumental punishment is also mOFC-dependent, as mOFC lesions prevented the extended retrieval of punishment contingencies relative to shams. In contrast, mOFC lesions did not prevent the re-acquisition of conditioned punishment that was learned prior to lesions being administered. Together, these results reveal that the mOFC does indeed regulate punishment learning and retrieval in a manner that is disassociated from any role in Pavlovian fear learning. These results imply that aberrant mOFC activity may contribute to the punishment insensitivity that is observed across multiple compulsive disorders.

---

There are relatively few studies of medial orbitofrontal cortex (OFC) function in rodents, despite its homologous region (i.e. mOFC, which can comprise anything from the more general ventral medial prefrontal cortex to the more specific Brodman’s Area 13) being of intense interest in human studies. This interest has arisen, in part, because aberrant activity in mOFC has been consistently identified in individuals with compulsive disorders, such as substance use disorder and obsessive-compulsive disorder (OCD) (Fineberg et al. 2011; Fettes et al. 2017; Moorman 2018). Thus, it is somewhat surprising that researchers have not made better use of rodent studies in order to make precise, causal inferences about the cognitive and behavioural consequences of dysregulated mOFC activity.

One prominent rodent study that has provided causal evidence of a link between aberrant mOFC activity and compulsive actions was that by Ahmari et al., (2013) who demonstrated that the aberrant excitation (via optogenetics) of mOFC terminals in ventral striatum can generate OCD-like grooming in mice. However, the nature of mOFC aberrance in compulsive individuals is complex, and is not limited to excitation but also features abnormal inhibition (Remijnse et al. 2006; Moorman 2018). Moreover, although compulsive-like grooming is a useful measure of repetitive behaviour (Kalueff et al. 2016), compulsive disorders are highly heterogeneous (Brady and Sinha 2005; Mataix-Cols et al. 2005) and grooming likely represents only a subset of related behaviours. Therefore, it is the aim of the current study to determine whether inactivating the mOFC produces another facet of compulsive disorders: insensitivity to punishment.

The inability to avoid punishing actions is a core characteristic of compulsive disorders (Hester et al. 2013; Figee et al. 2016). An individual with substance use disorder, for example, might persist in drug-seeking and drug-taking behaviours despite adverse effects on their relationships, health, and finances. Likewise, an individual with OCD might continue to wash their hands despite developing painful sores. Although such disorders are, of course, complex and multifaceted, and their neural underpinnings similarly intricate, ascertaining the causal consequences of aberrance in one part of this circuit (here, mOFC) for punishment sensitivity could provide insight into why mOFC activity is broadly aberrant across different compulsive disorders that share this core characteristic. Such a conclusion might be mitigated by the question of homology between human, primate, and rodent OFC, which remains somewhat contentious (Rolls and Grabenhorst 2008; Laubach et al. 2018). However, there are several recent circuit-based (Heilbronner et al. 2016) and function-based (Wallis 2012; Bradfield and Hart 2020) comparisons supporting the notion that there is at least some degree of functional, anatomical, and circuit-based homology between species.

Insight into why aberrant mOFC inhibition might lead to compulsive-like actions can be garnered from prior rat work (Bradfield et al. 2015; Bradfield et al. 2018). Specifically, we found that inactivating mOFC prevents rats from being able to infer the consequences of their actions, particularly when those outcomes are unobservable. Although all of the work done in support of this account was completed using appetitive outcomes, if mOFC also functions to infer the outcomes of actions when they are aversive it is straightforward to imagine that preventing this ability might also prevent individuals or animals from avoiding the actions that earn them. To be more specific, our prior work has shown that when a rat that presses a lever in order to receive a pellet it cannot see or smell, but must infer from prior lever-pressing experience, it can do so only with an intact mOFC. If the same is true of an animal who must infer an aversive outcome like footshock from their actions, in this case lever press, then it follows that a mOFC-inhibited animal will be unable to make that inference and will consequently fail to avoid that lever. This was the hypothesis that was tested in the current study. We further hypothesised, in line with our prior findings (Bradfield et al. 2015; Bradfield and Hart 2020), that this role for mOFC is specific to instrumental punishment and that its inactivation would not affect the acquisition or expression of Pavlovian conditioned fear. To test these hypotheses we compared animals with sham and excitotoxic lesions of the mOFC on several variants of a conditioned punishment procedure (Killcross et al. 1997), as this is a unique procedure which allows for a fully dissociated assessment of punishment-driven (instrumental) and fear-driven (Pavlovian) suppression of responding.

## Materials and Methods

### Experiment 1: Medial Orbitofrontal Cortex regulation of Conditioned Punishment Learning

#### Subjects

The subjects were experimentally naïve female and male Long-Evans rats (N = 32, female = 16, male = 16) supplied by the University of New South Wales (Sydney, NSW, Australia). Female rats weighed between 220-310 g, and male rats weighed between 390 - 530g at the beginning of the experiment. Rats were housed in home cages in groups of four in a temperature- and humidity-controlled room on a 12-hour light/dark cycle (lights on at 0700). Experiments were conducted during the light cycle. During behavioural training and testing, rats were maintained at ~85% of their free-feeding body weight and were allowed access to water *ad libitum*. All procedures were approved by the UNSW Animal Ethics Committee and are in accordance with the code outlined by the National Health and Medical Research Council (NHMRC) of Australia for the treatment of animals in research.

#### Apparatus

Behavioural procedures were conducted in standard operant chambers (Med Associates, Inc., East Fairfield, VT, USA), and controlled and recorded using Med-PC IV computer software (Med Associates, Inc.). A pellet magazine was base-centred on the right end wall, with one retractable lever on either side. Pellets (Bioserv, Biotechnologies) were delivered from a dispenser connected to the magazine. An infrared light situated at the magazine opening was used to detect head entries. A 10-second 3 kHz tone or 10-second 5 Hz flashing light were used as CSs, and 0.5-second footshocks with intensity ranging from 0.3 to 0.6 milliamps (mA) were delivered to the grid floor from a constant-current generator (see conditioned punishment procedure below for details). A house light was top-centred on the left end wall, and was illuminated throughout the entirety of each behavioural session.

#### Behavioural Procedures

##### Magazine training

Rats initially received one session of magazine training, during which both levers were retracted and pellets were delivered to the food magazine on a variable interval-60 (VI-60s) schedule. Magazine training was terminated and the rat removed from the operant chamber once either 20 pellets had been delivered or 30 minutes had elapsed, whichever came first.

##### Lever press training

Following magazine training, rats received 12 days of lever press training for pellets. On days 1-2, rats received two 30 min sessions, one on the left lever and one on the right (order counterbalanced). For these sessions, lever presses were continually reinforced (i.e. every lever press earned a pellet). After 20 pellets were delivered or 30 minutes had elapsed, whichever came first, the session was terminated, levers retracted, and house lights turned off. Lever training on days 3-6 was identical (one lever per day, order counterbalanced), except rats could earn as many pellets as the schedule allowed. Pellets were delivered on a variable interval schedule averaging 15sec (VI-15s) on days 3-4, which was increased to a VI-30s schedule on days 5-6.

Rats were then trained to press both levers (i.e. both were available throughout the entire session) on days 7-12 during 30 minute sessions. On day 7, responding on each lever was rewarded on a VI-15s schedule. On days 8-12, responding was reinforced on VI-30s schedule. Lever press training on days 8-11 was intended to equalise responding on each lever and remove any biases animals might show in responding on the left vs. right lever. To achieve this, a modified VI-30s schedule was implemented in which pellets were more frequently available on the non-preferred lever and less frequently available for delivery on the preferred lever (VI schedule adjusted as a ratio of responding on each lever). The last day (day 12) consisted of an unmodified VI-30s schedule on both levers to obtain an unbiased measure of pre-punishment lever pressing rates. Immediately after the last lever press training session, animals were returned to *ad libitum* access to chow in their home cages for 3 days prior to surgeries being carried out (see section on surgeries below for details).

##### Conditioned punishment

Following at least 7 days of recovery from surgeries, rats were again placed on food restriction as described. After 3 days of food restriction, rats received 5 days of conditioned punishment training. Each conditioned punishment training session lasted 60 mins during which both levers were presented throughout, and both continued to earn pellets on a VI-30s schedule as they had previously. In addition, the punished lever earned an aversive CS+ on a VI-60s schedule, whilst the unpunished lever earned a neutral CS- on a VI-60s schedule. The experiment was fully counterbalanced such that left lever was punished and the right lever was unpunished for half of the animals, whereas the other half received the opposite arrangement. For half of the animals in each arrangement, the tone was the CS+ and the flashing light was the CS-, and for the remaining half, the flashing light was the CS+ and the tone the CS-. The CS+ co-terminated in a 0.5 s footshock that increased in intensity over days. Daily foot-shock schedules over the 5 days were 0.3, 0.4, 0.4, 0.5, 0.5 mA for males, and 0.3, 0.3, 0.4, 0.4, 0.5 mA for females. The CS-terminated without consequence. If a lever press was scheduled to deliver both a pellet and a CS at the same time, only the CS was delivered due to its leaner schedule.

Punishment avoidance was determined as fewer lever presses on the punished versus the unpunished lever when no stimuli were present (i.e. during the inter-trial interval; ITI). Suppression of responding on both levers during the aversive CS+ was recorded and used to calculate a conditioned suppression ratio as a measure of Pavlovian fear (see statistical analysis for details).

##### VI-30 Conditioned Punishment

Following conditioned punishment training, rats completed another 5 days of conditioned punishment training with the reinforcement schedules altered such that the pellets and CSs were earned at double the frequency of initial punishment training. Specifically, pellet delivery was increased to a VI15s schedule (previously VI-30s) and CS presentation was increased to a VI30s schedule (previously VI60s).

#### Surgery

It should be noted that we initially proposed to investigate the differences between anterior and posterior lesions of the mOFC in conditioned punishment, although these groups were eventually collapsed across (see results). Thus, separate anterior and posterior co-ordinates are described.

Stereotaxic surgery was conducted under isoflurane anaesthesia (5% induction; 1– 2% maintenance). Each rat was placed in a stereotaxic frame (David Kopf Instruments, Tujunga, CA), after which they received a 0.1 ml subcutaneous injection of bupivacaine hydrochloride at the incision site. An incision was made into the scalp to expose the skull surface, and the incisor bar was adjusted to place bregma and lambda in the same horizontal plane. Anterior mOFC co-ordinates were (in mm relative to bregma): +5 anteroposterior, ±0.6 (males) or ±0.5 (females) mediolateral, and −4.6 dorsoventral. Posterior mOFC co-ordinates were (in mm relative to bregma): +3.8 anteroposterior, ±0.7 (males) or ±0.6 (females) mediolateral, −5.7 dorsoventral.

Excitotoxic lesions were made by infusing 0.3 μl of N-methyl-D-aspartate (NMDA: 10 mg/mL) in sterilised 0.1 M phosphate buffered saline (PBS) pH 7.2 over 3 min. The needle was left in place for 2 min prior to removal to allow for diffusion. Sham-operated rats underwent the same procedures but no excitotoxin was infused. Half of the sham lesions were performed at the anterior coordinates and the other half at the posterior coordinates. All rats received a subcutaneous injection of 0.1 ml carprofen and a 0.4 ml intraperitoneal injection of procaine penicillin solution (300 mg/kg). Rats were given 7 days to recover from surgery, after which they received 3 days of food deprivation prior to the commencement of experimentation. Rats were weighed and handled daily during this time.

#### Tissue preparation for histological analysis

Animals received a lethal dose of sodium pentobarbital (300mg/kg i.p., Virbac Pty. Ltd., Australia) 1-2 days following the completion of behavioural testing. Their brains were extracted and stored in a refrigerator kept at −30° Celsius. They were then frozen on Optimum Cutting Temperature (OCT) compound (Tissue-Tek, Sakura Finetek; Torrance, CA, USA) and coronally sectioned at 40μm through the mOFC using a cryostat (Leica CM1950, Leica Biosystems; Mt Waverley, Vic., Australia) maintained at approximately −18° Celsius.

Each section of the mOFC was collected directly onto a slide and was stained with NeuroTrace fluorescent nissl (Invitrogen, Eugene, OR, USA) and left to dry in darkness for at least 30 minutes. The slides were then cover-slipped with VectaShield (Vector Laboratories, Inc., Burlingame, CA, USA) and left to dry overnight in darkness. Sections were examined for lesion placements on a confocal microscope by a trained observer who was naïve to group allocation. Needle track marks were also examined for all groups including group SHAM.

#### Data analysis

Rates of responding per minute on the punished and unpunished levers during CS+, CS- and inter-trial interval (ITI; non-CS) periods were calculated. Punishment avoidance was measured as the rate of responding on the punished versus unpunished lever during the ITI. To determine if the punishment effect differed between groups, a 3-way, repeated measures ANOVA was conducted controlling the family-wise error rate at α = .05. For this ANOVA analysis, there was a between-subjects factor of group (i.e. Sham vs. Lesion) and a repeated measures factor of day (1-5) and of lever (punished vs. unpunished). Interactions with the ‘day’ factor were typically non-significant and thus not reported, although linear trend analyses were reported as evidence of learning in some instances. If an interaction (or interactions) were detected, then follow-up simple effects analyses were calculated to determine the source of the interaction.

Pavlovian conditioned fear was measured as the amount of suppression during the CS relative to ITI. Conditioned suppression ratios were calculated separately for the CS+ and CS-. This calculation was carried according to *Equation 1* for the CS+:

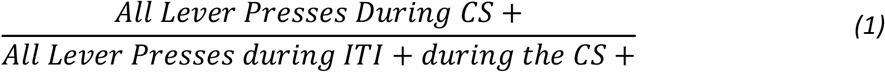

And according to *Equation 2* for the CS-:

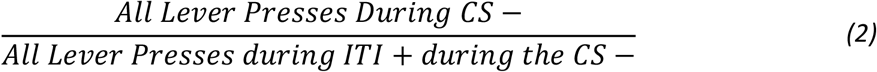

For these equations, a ratio below 0.5 indicates conditioned suppression (Pavlovian fear) to the CS, whereas a ratio of 0.5 or above indicates no fear. Once these values had been calculated separately for each CS, a 3 way repeated measures ANOVA, with a between-subjects factor of group, and a repeated measures factor of day (1-5) and another of CS (CS+ vs. CS-) controlling the familywise error rate at α = .05, was carried out to determine whether mean conditioned suppression ratios were higher to the CS+ than to the CS- for each group. Greater suppression to the CS+ relative to CS-indicates appropriate discrimination of Pavlovian fear.

### Experiment 2: Medial Orbitofrontal Regulation of Conditioned Punishment Retrieval

Experiment 2 was conducted identically to Experiment 1, except that mOFC lesions were administered post-punishment training, and animals were subject to retrieval tests. Specifically, lever press training took place as previously described, then rats received 7 days of pre-lesion conditioned punishment training. Following surgical procedures and recovery, rats received a 5-minute extinction test followed by a 10-minute test in which only pellets were delivered. Rats were then re-trained on conditioned punishment for 5 days and tested again in a 20 min test in which only pellets were delivered.

#### Subjects

Subjects were experimentally naïve female and male Long-Evans rats (N = 40, female = 20, male = 20) supplied by the University of New South Wales (Sydney, NSW, Australia). Female rats weighed between 220-310 g, and male rats weighed between 390 - 530g at the beginning of the experiment. Animals were housed and maintained in identical conditions to those described for Experiment 1.

#### Behavioural Procedures

All behavioural procedures were conducted as described for Experiment 1, except rats received one less day of the modified VI-30s schedule designed to equalise lever presses on each lever, which was sufficient for rats to show an unbiased response. Further, pre-surgery punishment training was completed for 7 rather than 5 days to ensure both groups had developed a strong punishment effect prior to surgery. The following tests were also carried out as described below.

##### 5 min extinction test and 10 min pellet-only test

Following recovery from surgery, rats received a 5 min extinction test in which both levers were extended, the houselight was turned on, but no outcomes (pellets, CSs ± footshock) were delivered. This test was immediately followed by a 10 min pellet-only test, in which both levers were extended and responding on them earned pellets on a VI-30s schedule as during lever press training, but no CSs ± footshock were delivered.

##### Conditioned Punishment Re-acquisition

Following these initial tests, rats received 5 days of re-training on conditioned punishment, which was conducted as previously described (pellets on VI-30s schedules, CSs ± footshocks on VI-60s schedules). The footshock intensity (mA) used for each rat was matched from its last day of pre-surgery punishment acquisition and kept consistent throughout re-training so as not to skew any behavioural effects of lesions.

##### 20 min pellet-only test

Following re-training, rats received another test in which only pellets were delivered on a VI-30s schedule. This time however, the pellet-only test was not preceded by an extinction test and lasted for 20 min.

*All surgical, histological, data collection and analyses procedures were conducted as described for Experiment 1*.

## Results

### Experiment 1: Medial Orbitofrontal cortex lesions impair conditioned punishment learning

Dysregulation of mOFC activity has been consistently identified in the brains of individuals with compulsive disorders such as OCD and substance use disorder (Maia et al. 2008; Fineberg et al. 2011; Moorman 2018). Insensitivity to punishment is characteristic of such disorders, yet causal evidence linking mOFC dysfunction to punishment insensitivity is lacking. Thus, we used a rodent model to examine whether lesions of mOFC caused insensitivity to punishment. To test this, we employed excitotoxic mOFC lesions in rats and assessed punishment avoidance and Pavlovian fear within a conditioned punishment procedure (Killcross et al. 1997).

In Experiment 1 we tested this hypothesis specifically with regards to conditioned punishment *learning*. Animals were first trained to press two levers (left and right) equally for pellets, after which they received sham or excitotoxic lesions of the mOFC. In the next stage of behavioural training, following recovery from surgeries, each lever continued to earn the pellets at the same rate (VI-30s), but each lever also began to earn a 10 sec conditional stimulus (CS). One CS (tone or flashing light) co-terminated with a mild footshock (CS+) while the alternate CS did not (CS-). This Pavlovian fear contingency causes the CS+ to acquire aversive value while the CS-remains neutral. Aside from pellets, pressing one lever occasionally yielded the CS+, an aversive consequence that instrumentally punished pressing of that lever; the other lever occasionally yielded the CS- and was thus unpunished.

Punishment learning was measured as avoidance of the punished lever relative to the unpunished lever during the inter-trial intervals (ITIs) between CS presentations, when rats’ choices were not influenced by CSs or footshocks. Rather, animals were required to infer from memory which lever would be punished and which would be unpunished. As the inference of unobservable outcomes is thought to rely on mOFC (Bradfield et al. 2015; Bradfield et al. 2018), we hypothesised that instrumental punishment learning would be impaired in group mOFC relative to group Sham. Pavlovian fear, on the other hand, elicits general decreases in reward-seeking (conditioned suppression), so was measured as suppression of lever-pressing during each CS relative to the ITI. The aversive CS+ was expected to produce conditioned suppression whereas the neutral CS- was not. Because Pavlovian fear is measured during CS presentations (i.e. when CS’s are observable) we hypothesised that it would not rely on mOFC and would be intact for both groups.

As previously noted, it was our initial intention to compare the relative roles of anterior mOFC and posterior mOFC (posterior mOFC is the anterior section of cingulate area 32V [AV32] according to Paxinos and Watson, 2014) in punishment learning, as we have previously found the anterior mOFC to be particularly important for the regulation of goal-directed action (Bradfield et al. 2018). However, for current experiments we were unable to separate these groups either anatomically or behaviourally, so they were collapsed across for all analyses.

#### Histology

Fig. 1A shows a sham anterior mOFC photomicrograph at +5.64 mm from bregma, and Fig. 1B show an excitotoxic lesions of anterior mOFC at +5.64 mm from bregma. Fig. 1C shows a sham posterior mOFC micrograph at +4 mm from bregma, and Fig. 1D shows an excitotoxic posterior mOFC lesion at +4 mm from bregma. Fig. 1E-F shows a representation of the overlapping lesion placements of all animals with anterior mOFC (Fig. 1E) and posterior mOFC (Fig 1F) lesions. If greater than 25% of the cell loss from a lesion extended more than a 1mm radius from the injection site, the rat was excluded from analysis. In total, 9 rats were excluded from Experiment 1 due to incorrect lesion placement or size, 1 rat died during surgery, and 1 rat was excluded due to a consistent and extreme preference for one lever (consistently > 100 more presses on one of the two levers) during lever press training, prior to surgeries. After exclusions, final numbers (*N* = 21) for groups in Experiment 1 were: sham control (*n* = 7), and mOFC lesions (*n* = 14: anterior *n* = 6, posterior *n* = 8). As mentioned, we collapsed across anterior and posterior mOFC lesion groups for analysis as they did not differ on any behavioural measure in either experiment (all p values > .05). Moreover, it is clear from Figure 1E-F that there was significant overlap between anterior and posterior lesions at approx. +4.68mm from bregma, such that we could not confidently separate these groups anatomically.

**Figure 1.**
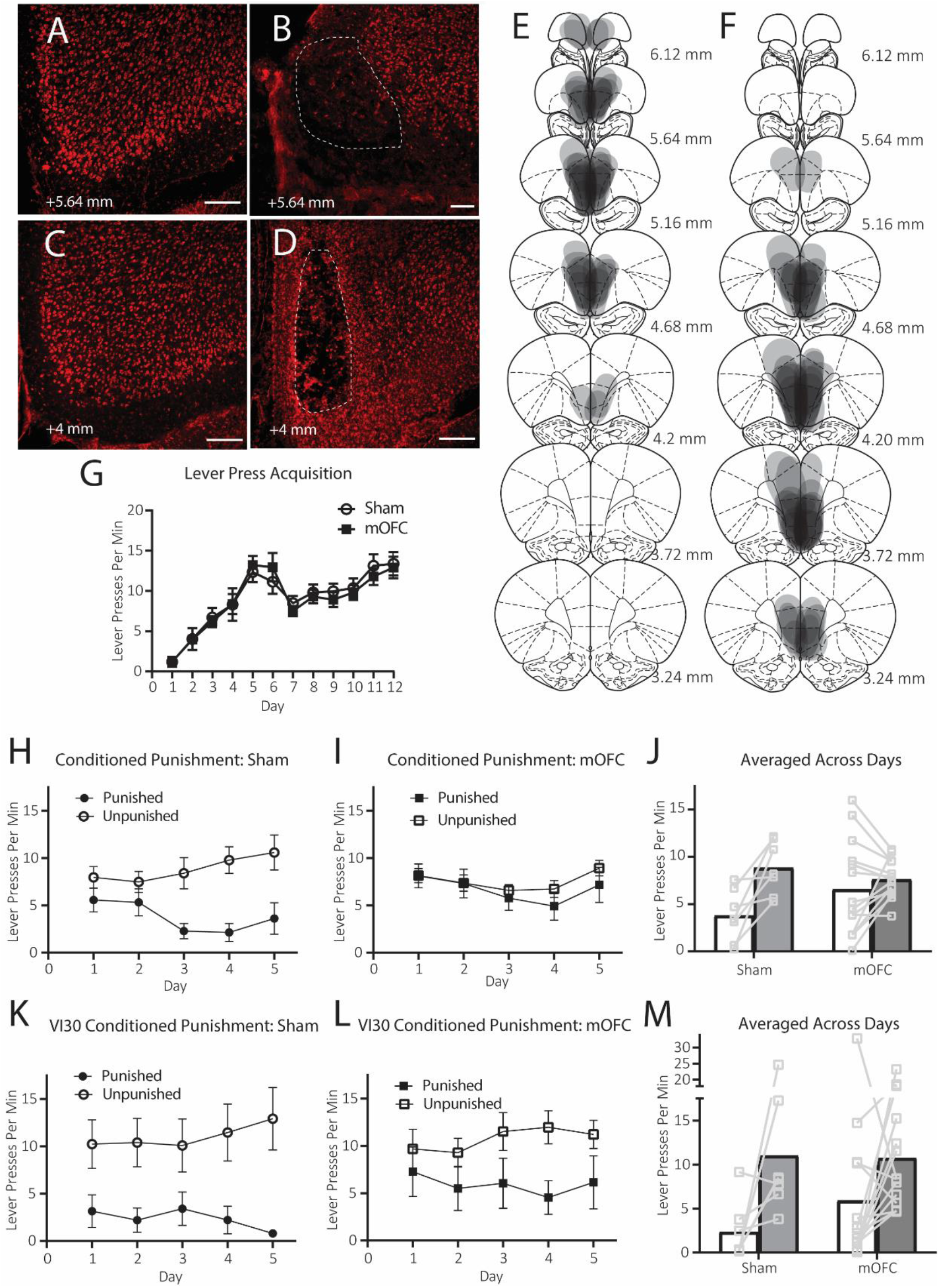
Excitotoxic lesions of medial orbitofrontal cortex prevent conditioned punishment learning. A-D) Representative photomicrographs of A) a sham anterior mOFC placement, B) an excitotoxic anterior mOFC lesion (both +5.64 mm from bregma, C) a sham posterior mOFC placement, and D) a posterior mOFC lesion (+4 mm from bregma). Scale bars = 100μm. F) Diagrammatic representation of anterior mOFC lesion placements, G) diagrammatic representation of posterior mOFC lesion placements, G) Mean (±1 SEM) lever presses per min during initial lever press acquisition, H) Mean (±1 SEM) lever presses per min during conditioned punishment training for group Sham, I) Mean (±1 SEM) lever presses per min during conditioned punishment training for group mOFC, J) Mean lever presses per min during conditioned punishment training for both groups, averaged over days, K) Mean (±1 SEM) lever presses per min during VI-30s conditioned punishment training for group Sham, L) Mean (±1 SEM) lever presses per min during VI-30S conditioned punishment training for group mOFC, M) Mean lever presses per min during VI-30s conditioned punishment training for both groups, averaged over days.

#### Behaviour Conditioned Punishment

##### Lever Press training

Lever press rates per min during initial lever press training are shown in Figure 1G (± SEM, averaged over levers). It is clear from this figure that groups Sham and mOFC both acquired lever press responding and did not differ in their acquisition (note that the dip in responding seen between Days 6-7 resulted from the increase in response competition after animals were switched from single-lever to double-lever protocols). Indeed, there was no main effect of group, F < 1, but there was a linear main effect of responding over days F (1,19) = 97.01, p = .00, that did not interact with group, F < 1.

##### Conditioned Punishment

The mean (±SEM) lever presses per min on the punished and unpunished levers over days during conditioned punishment training (during ITIs only) are shown in Figure 1H for group SHAM, in Figure 1I for group mOFC. The same data are shown with individual data points, averaged across the 5 days of conditioned punishment training for both groups in Figure 1J. From these figures, it is clear that group SHAM learned to avoid the punished lever during the ITI (unpunished > punished), but group mOFC did not (punished = unpunished). Statistical analysis showed that there was no main effect of group, F < 1, but there was a main effect of punishment F (1,19) = 11.33, p = .003, as well as a punishment × group interaction, F (1,19) = 5.023, p = .037. Follow up simple effects analysis reveal that this interaction comprised an intact punishment effect (unpunished > punished) for group SHAM, F (1,19) = 11.79, p = .003, but not for group mOFC (unpunished = punished), F < 1. Thus, this result demonstrates that mOFC lesions prevented animals from learning conditioned punishment.

##### Punishment Learning during VI-30s punishment

Results suggest that mOFC lesions produce a specific deficit in instrumental conditioned punishment learning. To test how pervasive this deficit might be, the same rats received 5 more days of conditioned punishment training, but we doubled the frequency of outcomes to determine whether this was sufficient to allow group mOFC to overcome the punishment learning deficit. That is, CSs and footshock were now delivered on a VI-30s schedule (increased from VI-60s) and pellets were now delivered on a VI-15s schedules (increased from VI-30s to keep pellet/CS ratio consistent).

Mean (±SEM) lever presses per min on the punished and unpunished levers over days during conditioned punishment (VI-30s) training (during ITIs only) are shown in Figure 1K for group SHAM and in Figure 1L for group mOFC. The same data are shown averaged across the 5 days of conditioned punishment (VI-30s) training for both groups in Figure 1M. It appears from these figures that the increase in reinforcement was sufficient for group mOFC to overcome their initial impairment because they, like group SHAM, responded selectively to the unpunished lever and avoided the punished lever. This is supported by statistical analysis because here was no main effect of group, F < 1, but there was a main effect of punishment F (1,19) = 7.54, p = .013, which this time, did not interact with group, F < 1.

#### Pavlovian Conditioned Fear

##### Conditioned Suppression during initial (VI-60s) Conditioned Punishment

Mean (±SEM) conditioned suppression ratios over days of conditioned punishment training are shown in Figure 2A for group SHAM and Figure 2B for group mOFC. It’s clear that animals in both groups SHAM and mOFC suppressed lever pressing during CS+ but not during CS-presentations. Statistically, there was no main effect of group, F < 1, but there was a main effect of CS, F (1,19) = 164.06, p = .00 that did not interact with group, F < 1. Thus, in contrast to instrumental punishment learning, mOFC lesions left Pavlovian fear conditioning to the CS+ intact.

**Figure 2.**
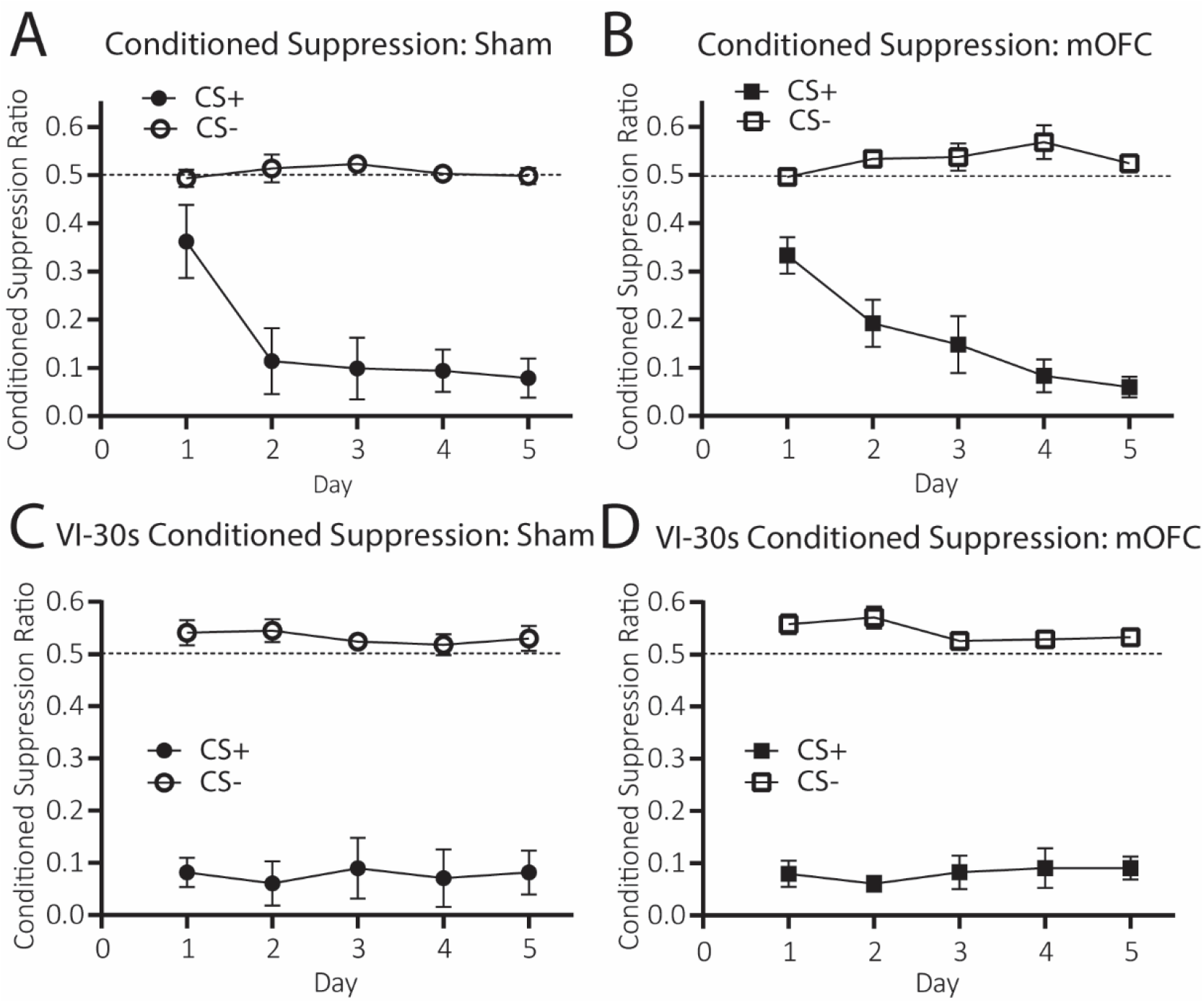
Excitotoxic lesions of medial orbitofrontal cortex leave Pavlovian fear conditioning intact. A) Mean (±1 SEM) conditioned suppression ratios to CS- and CS+ for group Sham during conditioned punishment training, B) Mean (±1 SEM) conditioned suppression ratios to CS- and CS+ for group mOFC during conditioned punishment training, C) Mean (±1 SEM) conditioned suppression ratios to CS- and CS+ for group Sham during VI-30s conditioned punishment training, B) Mean (±1 SEM) conditioned suppression ratios to CS- and CS+ for group mOFC during VI-30s conditioned punishment training.

##### Conditioned suppression during VI-30s punishment

Pavlovian fear was again unaffected during VI-30s punishment learning. Mean (±SEM) conditioned suppression ratios to the CS+ and CS-during VI-30s conditioned punishment training are shown over days in Figure 2C for group SHAM and Figure 2D for group mOFC. From these figures, it is clear that animals in both groups suppressed responding during CS+ but not CS-presentations. There was no main effect of group, F < 1, but there was a main effect of CS, F (1,19) = 379.72, p = .00 which did not interact with group, F < 1.

### Experiment 2: Medial orbitofrontal cortex lesions regulates the retrieval but not reacquisition of conditioned punishment

The results of Experiment 1 reveal a direct causal link between mOFC dysfunction and insensitivity to punishment. Sham rats learned to avoid a lever that earned a CS+ co-terminating in footshock, whereas rats with mOFC lesions were unable to adjust their behaviour to avoid punishment. This impairment was limited to instrumental punishment because mOFC lesions did not affect Pavlovian fear to the CS+. The fact that Pavlovian fear was intact suggests that mOFC lesions do not generally affect aversion or ability to suppress lever-pressing. Instead, it suggests that mOFC lesions induced a specific, punishment-insensitive phenotype identified in previous studies (Jean-Richard-dit-Bressel et al. 2019).

The effect of mOFC lesions in Experiment 1 was also limited to punishment learning, and whether mOFC is similarly necessary for the retrieval of punishment avoidance after it has been learned is unknown. This question is important because, as noted by Jean-Richard-Dit-Bressel, Killcross, and McNally (2018), in real-world situations punishment is rarely immediate (e.g. drug withdrawals occurring the following day, financial consequences might occur months later). Thus, in order for individuals to successfully avoid punishments that may not occur until later, they must retrieve previously learned punishment contingencies when no punishing outcomes are currently being presented.

To determine whether mOFC is necessary for the retrieval of punishment avoidance after it has been learned, in this experiment we lesioned the mOFC after conditioned punishment training and gave animals retrieval tests in which no punishing outcomes were delivered. Specifically, following recovery from surgery, animals received a 5 min extinction test (i.e. no outcomes delivered) followed by a 10 min test in which only pellets were delivered. Because no CSs or footshocks were delivered on either test, responding in accordance with punishment contingencies required these outcomes to be inferred from memory. Thus group mOFC were expected to be impaired (unpunished = punished) on this test relative to group Sham (unpunished > punished). Following testing, we retrained animals on conditioned punishment for 5 days and then tested their retrieval again in a 20 min pellet-only test.

#### Histology

Figure 3A-B shows a representation of the overlapping lesion placements of all animals with anterior mOFC (Figure 3A) and posterior mOFC (Figure 3B). If greater than 25% of the cell loss from a lesion extended more than a 1mm radius from the injection site, the rat was excluded from analysis. Five rats were excluded from Experiment 2 due to incorrect lesion placement or size, 3 rats died during surgery, 6 rats were excluded due to malfunctioning operant chambers, and 1 rat was excluded due to a consistent and extreme affinity for the punished lever during conditioned punishment training prior to surgery (i.e. > 100 more presses on punished than unpunished lever every day). After exclusions, final numbers (*N* = 25) for groups in Experiment 2 were: group SHAM (*n* = 11), and mOFC lesions (*N* = 14: anterior *n* = 8, posterior *n* = 6). As in Experiment 1, we collapsed anterior and posterior mOFC lesion groups for analysis as and there was significant overlap of lesions, particularly at +4.68 mm anterior to bregma, and they did not differ significantly on any measure (all p values > .05).

**Figure 3.**
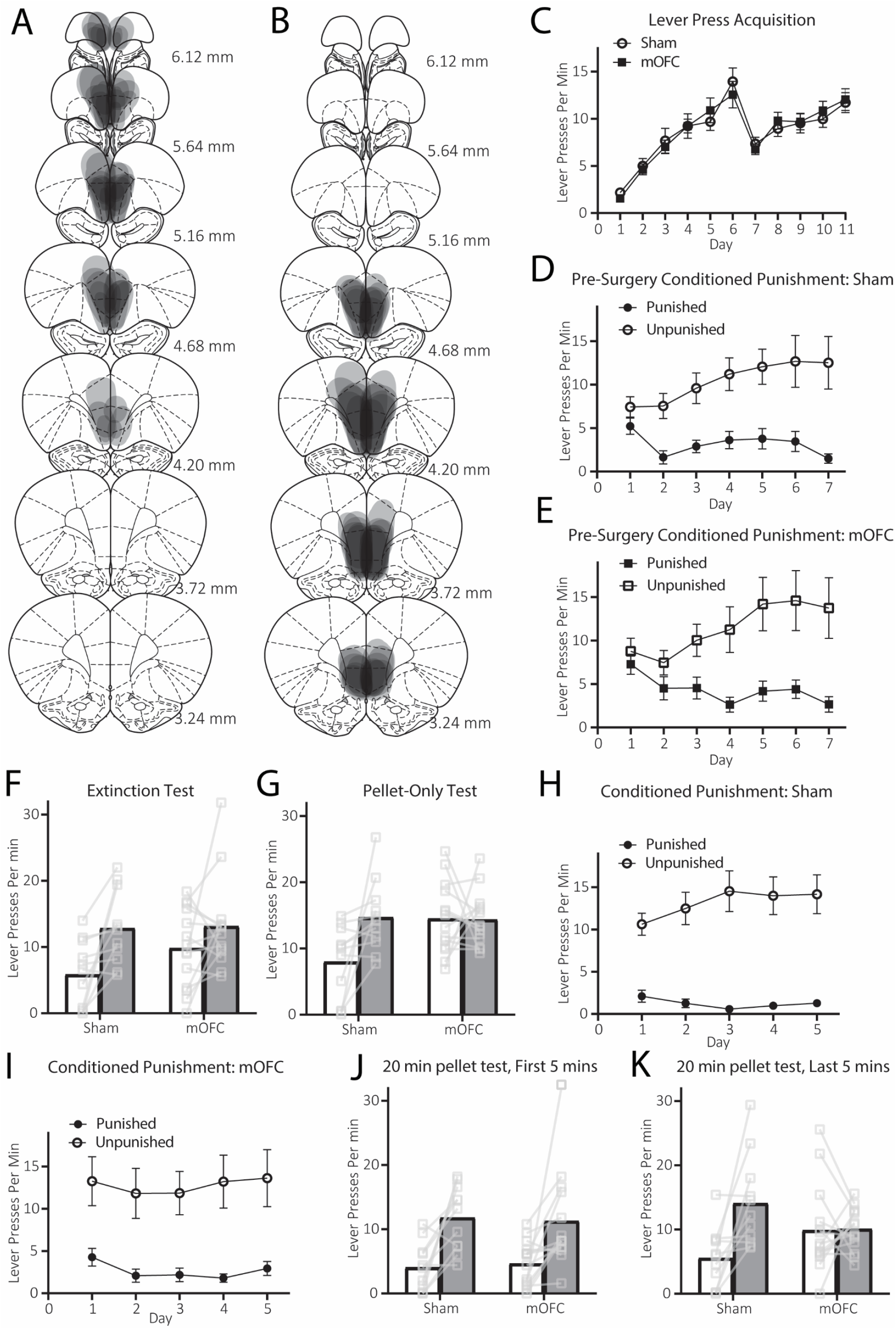
Excitotoxic lesions of medial orbitofrontal cortex impair conditioned punishment retrieval. A) Diagrammatic representation of anterior lesion placements, B) diagrammatic representation of posterior lesion placements, C) Mean (±1 SEM) lever presses per min during initial lever press acquisition, D) Mean (±1 SEM) lever presses per min during pre-surgery conditioned punishment training for group Sham, E) Mean (±1 SEM) lever presses per min during pre-surgery conditioned punishment training for group mOFC, F) Mean lever presses per min for both groups during the 5 min extinction retrieval test, G) Mean lever presses per min for both groups during the 10 min pellet-only retrieval test, H) Mean (±1 SEM) lever presses per min during conditioned punishment retraining for group Sham, I) Mean (±1 SEM) lever presses per min during conditioned punishment retraining for group mOFC, J) Mean lever presses per min during the first 5 mins of the 20 min pellet-only retrieval test, K) Mean lever presses per min during the last 5 mins of the 20 min pellet-only retrieval test.

#### Behaviour Conditioned Punishment

##### Lever Press training

Lever press rates per min during initial lever press training are shown in Figure 3C (± SEM, averaged over levers) shown with data split according to the groups assigned at surgery. This figure shows that the groups SHAM and mOFC both acquired lever press responding and did not differ in their acquisition (again there is a dip in responding from Days 6-7 due to increased response competition that occurred as animals were transferred from the single-lever to the double-lever protocol). There was no main effect of group, F < 1, but there was a linear main effect of responding over days (1,19) = 141.714, p = .00, that did not interact with group, F < 1.

##### Conditioned Punishment (Pre-Surgery)

Conditioned punishment training occurred across 7 days prior to lesion surgeries. Data is shown according to the groups assigned during surgery. Mean (± SEM) rates of responding on the punished and unpunished levers over days (during ITIs only) are shown in Figure 3D for group SHAM and Figure 3E for group mOFC. It is clear that both groups SHAM and mOFC acquired punishment avoidance prior to lesion surgeries and did not differ in their acquisition, as both responded more on the unpunished relative than the punished lever. There was no main effect of group, F < 1, but there was a main effect of punishment, F (1, 23) = 14.97, p = .001, that did not interact with group, F < 1.

##### Extinction test

Mean (± SEM) lever presses for groups SHAM and mOFC during the 5 min extinction test are shown in Figure 3F. This figure shows that group SHAM responded according to the punishment contingencies (unpunished > punished). Although this effect appears to be somewhat attenuated in group mOFC, no statistical differences between the two groups were detected. Specifically, there was no main effect of group, F (1,23) = 3.112, p = .091, but there was a main effect of punishment, F (1,23) = 11.435, p = 0.003, that did not interact with group, F (1,23) = 1.4, p = .25. These findings suggest that both sham and mOFC-lesioned animals can retain and express the punishment contingencies they learned prior to the mOFC lesion, at least upon initial testing in extinction.

##### Pellet-only Test

Mean (± SEM) lever presses for groups SHAM and mOFC during the 10 min pellet-only test are shown in Figure 3G. This figures shows that group SHAM continued to respond in accordance with the punishment contingencies (unpunished > punished) but group mOFC did not (unpunished = punished). This is supported by statistical analysis, as there was a main effect of group, F (1,23) = 6.13, p = .021, indicating that group mOFC responded more overall than did group SHAM, a marginal main effect of punishment, F (1,23) = 3.97, p = .058, but there was a significant punishment × group interaction, F (1,23) = 4.34, p = .049. Follow-up simple effects analysis showed this interaction consisted of a significant effect of punishment in group SHAM, F (1,23) = 7.41, p = .012, but not group mOFC, F < 1. This result suggests that animals in group SHAM continued to apply the punishment contingencies they had learned pre-surgery (i.e. unpunished > punished) throughout the test, despite the availability of pellets and the absence of CSs and footshock. This is consistent with previous results demonstrated by Jean-Richard-Dit-Bressel and McNally (2016) in which control animals continued to apply (primary) punishment contingencies throughout the entirety of a 30 min pellet-only test. In contrast, punishment retrieval was impaired for group mOFC who responded equally on both levers.

Why mOFC lesions impaired performance on this test but not the prior extinction test is unclear. Indeed, we had predicted that mOFC lesions would impair performance on both tests due to the absence of punishing outcomes. One possibility is that group mOFC’s performance *was* impaired on both tests, but only the pellet test was sufficiently powered to detect a group × punishment interaction, perhaps due to more consistent responding across the 10 mins which reduced variability. Another possibility is that the switch in contingencies from extinction to pellets caused confusion or interference in group mOFC that was not present in sham animals. A final possibility is that both groups initially recalled the punishment contingencies, but that their extinction was facilitated in group mOFC relative to group SHAM.

To distinguish between these possibilities, we re-trained animals on the conditioned punishment contingencies for 5 days then gave them another pellet-only test, only this time the pellet-only test lasted for 20 min and was not preceded by any period of extinction. Sham animals were once again expected to maintain responding in accordance with the punishment contingencies for the entire duration of the test. If mOFC lesions produce a general decrement in the retrieval of punishment contingencies which the prior extinction test was underpowered to detect, group mOFC should be impaired for the entire 20 mins. Alternatively, if group mOFC were previously impaired due to the contingency change from extinction to pellets, then group mOFC should display intact performance for the entire 20 mins, as no contingency change is applied on this test. Finally, if extinction of the punishment contingencies was facilitated in group mOFC relative to group Sham, on the 20 min test this group should once again initially respond according to these contingencies (unpunished > punished) then revert to equal lever pressing on both levers (unpunished = punished) for the latter part of the test.

##### Conditioned punishment re-training

Mean (± SEM) lever presses per min during the reacquisition of conditioned punishment is shown over days in Figure 3H for group SHAM, in Figure 3I for group mOFC. It is clear from these figures that punishment (unpunished > punished) is intact for both groups during retraining. Statistical analysis supports this conclusion because there is no main effect of group, F < 1, but there is a main effect of punishment, F (1,23) = 28.99, p = .00, and no punishment × group interaction, F < 1. This suggests that, in contrast to initial conditioned punishment learning in Experiment 1, mOFC lesions do not produce a decrement in the re-acquisition of conditioned punishment.

##### 20 min pellet-only test

The question of interest was whether group mOFC would respond in accordance with the punishment contingencies (unpunished > punished) for none of the test, some of the test, or all of the test. We therefore split the data from the 20 min pellet test into four 5 min bins and examined the first 5 min and the last 5 min separately (note that we also split the data into two 10 min bins and compared the first 10 vs. the last 10 min and found an identical pattern of responding and statistical outcome, but the effects were slightly clearer when the first 5 and last 5 mins are compared).

Mean (± SEM) lever presses during the first 5 min of the 20 min pellet-only test are shown in Figure 3J. This figure clearly shows that in the first 5 mins, both groups SHAM and mOFC responded in accordance with the punishment contingencies (unpunished > punished). Statistically, there was no main effect of group, F < 1, but there was a main effect of punishment, F (1,23) = 21.63, p = .00, which did not interact with group, F < 1.

Mean (± SEM) lever presses during the last 5 min of the 20 min pellet-only test are shown in Figure 3K. It is clear from this figure that, in contrast to responding during the first 5 mins of this test, performance was impaired for group mOFC (unpunished = punished) relative to group Sham (unpunished > punished). This is supported by statistical analysis, because although there was no main effect of group, F < 1, and there was a main effect of punishment, F (1,23) = 7.98, p = .01, and there was also a group × punishment interaction, F (1,23) = 7.27, p = .013. Follow up simple effects reveal that this interaction consisted of a significant effect of punishment (unpunished > punished) for group SHAM, F (1,23) = 13.62, p = .001, but not group mOFC (unpunished = punished), F < 1.

This result suggests that the prior test results were not an artefact of an underpowered test, or the contingency change from extinction to pellets-only. Rather, animals with mOFC lesions rapidly lost punishment avoidance across these punishment-extinction tests, suggesting that extinction of the punishment contingencies was facilitated for group mOFC relative to group Sham. Thus, these results present only partial support for our prediction that mOFC lesions would prevent the retrieval of punishment contingencies. Reasons for this divergence from our hypothesis are discussed below.

#### Pavlovian Conditioned Fear

##### Conditioned Suppression (during pre-surgery conditioned punishment)

Mean (± SEM) conditioned suppression ratios to the CS+ and CS-over days of pre-surgery conditioned punishment training are shown in Figure 4A for group SHAM and Figure 4B for group mOFC. It is clear from these figures that animals in both groups acquired specific fear to the CS+ and not the CS- and did not differ in their acquisition. This is confirmed by statistical analysis as there was no main effect of group, F < 1, a main effect of CS, F (1,23) = 267.75, p < .001, that did not interact with group, F < 1.

**Figure 4.**
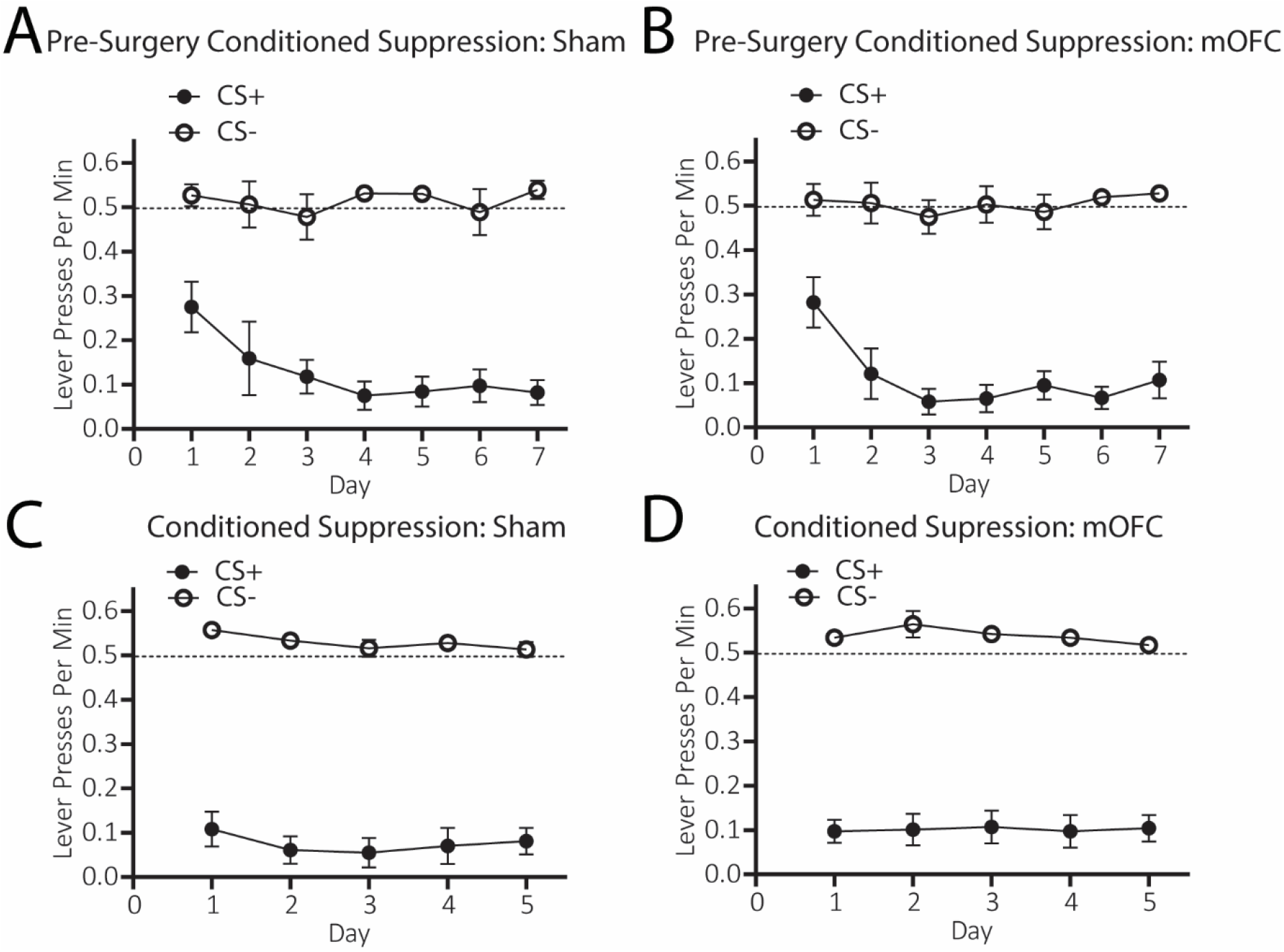
Excitotoxic lesions of medial orbitofrontal cortex leave Pavlovian fear conditioning intact. A) Mean (±1 SEM) conditioned suppression ratios to CS- and CS+ for group Sham during pre-surgery conditioned punishment training, B) Mean (±1 SEM) conditioned suppression ratios to CS- and CS+ for group mOFC during pre-surgery conditioned punishment training, C) Mean (±1 SEM) conditioned suppression ratios to CS- and CS+ for group Sham during conditioned punishment retraining, D) Mean (±1 SEM) conditioned suppression ratios to CS- and CS+ for group mOFC during conditioned punishment retraining.

##### Conditioned Suppression during conditioned punishment re-training

Mean (±SEM) conditioned suppression ratios to the CS+ and CS-during conditioned punishment re-training are shown in Figure 4C for group SHAM and Figure 4D for group mOFC. It is clear that both groups once again showed selective conditioned suppression to the CS+ than to the CS-, indicating intact Pavlovian fear for both groups. Statistical analysis reveals that there was no main effect of group, F < 1, but there was a main effect of CS, F (1, 23) = 387.63, p =< .001 that did not interact with group, F < 1.

## Discussion

Together, current results demonstrate a causal role for mOFC in instrumental punishment avoidance, but not Pavlovian conditioned fear. In Experiment 1, mOFC lesions prevented conditioned punishment learning. Sham controls demonstrated intact conditioned punishment learning by avoiding a punished lever that earned a CS+ and footshock whilst continuing to respond on an unpunished lever that earned a neutral CS-, whereas animals with mOFC lesions responded equally on punished and unpunished levers. This deficit was specific to instrumental punishment learning because Sham and mOFC groups both selectively suppressed lever-pressing during CS+ relative to CS-presentations. This previously-identified phenotype of punishment insensitivity has been attributed to failures in detecting lean punishment contingencies (Jean-Richard-dit-Bressel et al. 2019). Consistent with this, we found that doubling the frequency of outcomes (i.e. pellets, CS+, CS-, and footshock) allowed group mOFC to overcome their initial impairments and avoid the punished response in favour of the unpunished response.

Experiment 2 tested whether mOFC lesions prevented the retrieval as well as the learning of conditioned punishment contingencies. For this experiment, lesions were administered after conditioned punishment training. Following recovery, animals were given a 5 min extinction test (i.e. no outcomes delivered) followed by a 10 min test in which both levers earned only pellets. In the 5 min extinction test, both groups demonstrated statistically intact punishment avoidance. However, only Sham group continued to respond in this manner in the subsequent pellet-only test, whereas mOFC-lesioned animals responded indiscriminately on both levers (i.e. unpunished = punished). Rats were subsequently re-trained on conditioned punishment for 5 days, and then given a pellet-only test for 20 min. During retraining, both groups Sham and mOFC were able to selectively respond to the unpunished lever over the punished lever, suggesting that mOFC-lesioned rats could relearn punishment contingencies. On the final pellet-only test, both groups once again initially responded according to the punishment contingencies (unpunished > punished), but only Sham animals did so throughout the entire test. Group mOFC, on the other hand, responded equally on both levers for the latter half of the test, replicating the pattern observed in prior tests. This suggests mOFC lesions may accelerate punishment extinction.

Together, these findings provide partial support for our hypothesis that mOFC lesions would prevent animals from inferring aversive outcomes as the consequence of their actions. The results of Experiment 1 support this account because mOFC lesions initially prevented conditioned punishment learning as predicted, which is measured during the ITI when CSs and punishing footshocks had to be inferred. Moreover, when outcomes were doubled in frequency and thus made more ‘observable’, mOFC-lesioned animals were able to overcome their initial impairment. The results of Experiment 2, however, are more difficult to reconcile with this account. During both sets of retrieval tests in Experiment 2, it appeared that group mOFC did initially retain and express the punishment contingencies (i.e. unpunished > punished), despite extinguishing more quickly than for group Sham. Furthermore, re-acquisition of conditioned punishment after mOFC lesion surgery was completely intact for group mOFC, even though punishment was again measured during the ITI when outcomes had to be inferred. These effects suggest that, under some circumstances at least, group mOFC were able to infer punishing outcomes even when unobservable. Therefore, taken together with our prior findings (Bradfield et al. 2015; Bradfield et al. 2018), these results suggest that the role for mOFC in inferring the unobservable outcomes of actions might be limited to appetitive outcomes. This would imply that mOFC regulates instrumental aversive learning via a different psychological mechanism. Although the identity of that mechanism is not immediately clear, what is apparent from current results is that the mOFC plays a conditional role in punishment learning and retrieval, and that it is particularly important for these processes when the presentation of punishing outcomes is sparse. This role for mOFC could be viewed as consistent with its proposed role in probability estimation (Stopper et al. 2014).

There is another possibility, however, that could reconcile our initial interpretation of mOFC function with current results. It has been noted that with extended training, animals can learn to avoid a punished action out of habit (Ostlund and Balleine 2008). Thus it is possible, perhaps even likely, that during the 7 days of pre-surgery conditioned punishment training in Experiment 2, animals learned to avoid the punished lever in a habitual manner. Our account clearly predicts that habit learning should be unaffected by mOFC lesions because the ability to infer outcomes does not contribute to habitual stimulus-response (S-R) associations. Punishment retraining in Experiment 2 could have simply reinstated this habit in group mOFC, causing their re-acquisition to be intact relative to group SHAM. During retrieval tests, this habit (unpunished > punished) could likewise have been applied for both groups initially, but when this habit ‘failed’ in the absence of any punishing outcomes, animals returned to goal-directed responding as is known to occur under extinction conditions (Dezfouli et al. 2014). For group Sham, this meant continued avoidance of the punished lever based on the goal-directed inference that presses could earn the unobservable CS+ and footshock. For group mOFC however, who could not infer these outcomes, goal-directed responding would have been elicited on the basis of the observable pellet outcomes only, which were earned equally by each lever (unpunished = punished).

One problem for this account is why group mOFC were able to use habitual S-R associations to re-acquire conditioned punishment in Experiment 2, but were unable to use these associations to acquire conditioned punishment in Experiment 1. Because S-R learning is thought to update more slowly than flexible, goal-directed learning (Dickinson and Balleine 1995), and punishment training was conducted for longer in Experiment 2 than Experiment 1 (5 days of punishment in Experiment 1 vs. 7 days of pre-surgery punishment training in Experiment 2), it was perhaps the extra training sessions in the latter experiment over which the habit was critically formed. Indeed, there is some evidence that habitual S-R punishment avoidance had begun to occur even by day 3 of conditioned punishment in Experiment 1, as a small unpunished > punished difference appears to emerge in group mOFC’s performance from day 3 onwards (see Figure 1I). It is possible that group mOFC would have continued to develop a habitual avoidance of the punished lever in this experiment even without doubling the frequency of outcomes after 5 days of training.

Another surprising finding was the lack of any behavioural differences in animals that received anterior versus posterior mOFC lesions. As stated, we have found in previous studies (Bradfield et al. 2018) the anterior but not posterior mOFC to be particularly important for the inference of outcome representations. Here, we targeted each sub-structure separately to assess whether they differentially regulate punishment learning and expression, but did not find any support for this hypothesis. As we were unable to separate these lesions anatomically, due to a significant site of overlap (centred at +4.68mm anterior to bregma), it is possible that all of our current lesions simply targeted the same functionally important mOFC region. However, it is also possible that posterior mOFC may not contribute to the inference of appetitive outcomes, but does contribute to punishment sensitivity. In fact, if this were the case it would be consistent with several findings (Gourley et al. 2010; Münster and Hauber 2018) in which inactivation of [posterior] mOFC produces more persistent lever-press responding than in controls, and a higher ‘break point’ at which animals stop responding for progressively more sparse outcomes. This ever-increasing scarcity of outcomes could be interpreted as a ‘punishment’ of lever presses in this task, and insensitivity to this punishment as a result of posterior mOFC inactivation may have contributed to the observed increase in breakpoint.

Regardless of the anatomical specificity, current results clearly demonstrate a causal link between dysregulation of mOFC function and sensitivity to punishment in a manner that cannot be attributed to differences in Pavlovian fear learning. This is important in ruling out current effects being a result of overlap with anatomically adjacent prefrontal cortical regions, such as prelimbic and infralimbic cortex (more recently referred to as part of cingulate areas A32V and A25, respectively; Laubach et al. 2018; Paxinos and Watson 2014), which are known to have a role in Pavlovian fear. That is, prelimbic cortex has been shown to regulate Pavlovian fear expression directly (Quirk et al. 2000; Laurent and Westbrook 2009) and our lesions did not affect Pavlovian fear expression at any stage. Infralimbic cortex does not affect fear expression, but it does affect fear extinction (Quirk et al. 2000; Milad and Quirk 2002). Although there was no possibility of direct fear extinction to the CS+ in current experiments, because the CS+ was never presented in the absence of footshock, it is possible that indirect fear extinction occurred to the lever-associated stimuli. That is, as animals successfully learn to avoid the punished lever, they experience fewer punishing outcomes despite the continued presence of lever-associated stimuli, essentially creating an extinction contingency. Had lesions prevented this extinction, we would expect to see facilitated punishment learning (if anything) because lesioned animals would continue to be afraid of the punished lever stimuli, whereas fear of these stimuli would have reduced (extinguished) in sham animals. As this was clearly not the observed result, and as prelimbic and infralimbic lesions have also been shown to play no role in primary punishment paradigms (Jean-Richard-Dit-Bressel and McNally 2016), it is unlikely that current effects could have been caused by lesion overlap with either prelimbic or infralimbic cortex.

Although there is still much to be determined regarding mOFC’s regulation of punishment learning and retrieval, the discovery of a causal link is an important starting point, and one that has multiple implications. For instance, given the strong reciprocal links between mOFC and the basolateral amygdala (Hoover and Vertes 2011; Malvaez et al. 2019), and the central role of basolateral amygdala in punishment avoidance (Killcross et al. 1997), the mOFC-basolateral amygdala pathway is a prime candidate for the neural circuitry that underpins conditioned punishment. Medial OFC also has a number of other key connections that could mediate punishment learning, such as its strong outputs to nucleus accumbens core (Hoover and Vertes 2011) which is central to motivated behaviour (Corbit et al. 2001; Corbit and Balleine 2011). Moreover, it has now been demonstrated several times that the orbitofrontal cortex accommodates a functionally heterogeneous population of neuronal ensembles (Schoenbaum et al. 1999; Tremblay and Schultz 1999; Padoa-Schioppa and Assad 2006), some of which have been individually manipulated and demonstrated to regulate unique types of behaviour (Jennings et al. 2019). Although most of these studies have focused on lateral OFC, it is possible that heterogeneous neuronal ensembles also exist within mOFC, and that their coordination is what allows mOFC to form complex, abstract outcome representations regardless of appetitive or aversive valence. The non-specific excitotoxic lesions in the current study would have indiscriminately targeted all of these ensembles, but with the application of the increasingly specialised tools, future studies can begin to unravel the potential contributions of each with great precision. Furthermore, as lateral OFC inactivation has also been shown to affect punishment (Orsini et al. 2015; Jean-Richard-Dit-Bressel and McNally 2016; Verharen et al. 2019) similar studies of the lateral OFC’s contributions to conditioned punishment learning and expression will also be of great interest to future studies.

In conclusion, current results demonstrate a causal relationship between dysregulation of mOFC function and insensitivity to punishment but not Pavlovian fear learning. Further, mOFC activity appears to be particularly important for punishment learning when aversive outcomes are less frequently observable, and when experience with punishment is not sufficient to cause habitual avoidance of the punished action. This could have myriad interesting clinical implications if translatable. For instance, individuals suffering from compulsivity and insensitivity to punishment due to mOFC dysfunction might reinstate their sensitivity if punishing outcomes were made more observable or consistent.

## Acknowledgements

We thank Gavan McNally for helpful discussions regarding these data.

